# Whole genome sequencing analysis identifies rare, large-effect non-coding variants and regions associated with circulating protein levels

**DOI:** 10.1101/2023.11.04.565589

**Authors:** Gareth Hawkes, Kartik Chundru, Leigh Jackson, Kashyap A. Patel, Anna Murray, Andrew R Wood, Caroline F Wright, Michael N Weedon, Timothy M Frayling, Robin N Beaumont

**Affiliations:** Department of Clinical and Biomedical Sciences, Faculty of Health and Life Sciences, University of Exeter, Exeter, EX1 2LU; Faculty of Medicine, Department of Genetic Medicine and Development, CMU,1 rue Michel-Servet, Geneva, Switzerland

## Abstract

The role of non-coding rare variation in common phenotypes is largely unknown, due to a lack of whole-genome sequence data, and the difficulty of categorising non-coding variants into biologically meaningful regulatory units. To begin addressing these challenges, we performed a *cis* association analysis using whole-genome sequence data, consisting of 391 million variants and 1,450 circulating protein levels in ∼20,000 UK Biobank participants. We identified 777 independent rare non-coding single variants associated with circulating protein levels (*P*<1×10^-9^), after conditioning on protein-coding and common associated variants. Rare non-coding aggregate testing identified 108 conditionally independent regulatory regions. Unlike protein-coding variation, rare non-coding genetic variation was almost as likely to increase as decrease protein levels. The regions we identified overlapped predicted tissue-specific enhancers more than promoters, suggesting they represent tissue-specific regulatory regions. Our results have important implications for the identification, and role, of rare non-coding variation associated with common human phenotypes.

## Introduction

Rare genetic variants in non-coding regions of the human genome can cause severe rare disease^1,2^, but their role in common, complex traits is still largely unknown. Array-based genome-wide association studies (GWAS), which use imputed genotype measures, have identified tens of thousands of common variant associations with human disease^3^, the majority of which are located outside of coding regions^4^. However, efforts to identify rare variants associated with common phenotypes have been largely limited to the coding regions, using exome sequencing data. This limited progress is due to the lack of whole-genome sequence data in large studies and the difficulty of defining biologically meaningful non-coding regulatory genomic units.

The analysis of whole-genome sequencing (WGS) data could provide important insight into relevant genes and their regulation that complements the knowledge gained from exome sequencing and array-based studies. WGS allows us to examine the role of intronic, proximal and distal regulatory elements, and covers the entire allele frequency spectrum in a population, including a very large proportion of variants that are observed only once or twice even in a very large sample.

The identification of non-coding regulatory elements could provide important insight into the tissue-specific roles on the regulation of nearby genes. There is considerable evidence that the non-coding genome is functionally important. For example, based on population genetic metrics such as constraint, the amount of non-coding DNA that is functional could be 4-5x greater than the amount of coding sequence^5^, and 10% and 6% of promoters and enhancers respectively are under as much mutational constraint as coding regions^6^. Enhancers have also been shown to be more tissue-specific than promoters^7^.

There are very few studies of WGS data in the context of common phenotypes. Recent examples from TOPMed^8^ have considered lipid-levels^9,10^ (N = 66,000) and blood pressure^11^ (N = 51,456) but found few novel signals, possibly because of the complexity of the phenotypes and relatively small sample sizes for the detection of novel disease-associated rare variants.

The UK Biobank’s (UKB) release of circulating protein data, in combination with WGS, provides an unprecedented opportunity to test the impact of rare non-coding genetic variation on common, biologically proximal, and well-measured human phenotypes. Three recently published studies^12–14^ described the 2023 release of this data on up to 2,923 circulating proteins in 54,306 individuals. These studies focused on conventional array-based GWAS approaches or exome sequencing, analysing between 0.5 million and 58 million variants, but did not attempt to use the full range of allelic variation consisting of >700 million variants available in the WGS data. These studies identified a large number of pQTLs (protein Quantitative Trait Loci) including with rarer single variants and coding variants. Firstly, Eldjarn et. al (2023)^12^ identified 30,062 pQTLs in a single-variant analysis of genetic data imputed from the UKB 150,119 whole-genome sequences with 2,931 measured protein levels and compared results with proteomics derived from an Icelandic cohort with whole-genome sequences. Secondly, Dhindsa *et al.* (2023)^13^ identified 5,433 pQTLs in an exome-sequencing analysis, and performed aggregate testing within the coding regions, identifying 1,962 gene-protein associations. Finally, Sun *et al* (2023)^14^ identified 14,287 pQTL single-variants using a conventional GWAS of array-based imputed data.

Using WGS data and circulating protein levels as exemplar traits, we tested two related hypotheses: 1) non-coding single variants, not currently detectable by GWAS array or exome sequencing, contribute to common human phenotypes with similar effects to coding variants, and 2) we can identify aggregates (groups) of rare non-coding genetic variation in regulatory regions of the genome associated with human phenotypes, akin to gene-level collapsing analyses in exome sequences. Importantly, and in contrast to the previous three papers on the UKB proteomic data, we used the full range of DNA sequence variation detected with short read WGS, providing information on ∼400 million variants, but limited the search for each phenotype to *cis* regions around the protein-coding gene from which the protein derived.

## Methods Summary

We performed primary discovery association analyses for 1,450 measured circulating protein levels using annotated WGS data on 20,038 individuals of inferred European genetic ancestry from the UKB, a population cohort from the United Kingdom. The vast majority (95.6%) of samples were of genetically-inferred European genetic ancestry. Thirteen proteins that were either fusion proteins or did not directly match to an HGNC gene symbol were excluded (**ST1**). For each measured protein, we performed both single variant (minor allele count (MAC) ≥ 5) and genomic aggregate association tests (minor allele frequency (MAF) < 0.1%) in a *cis-*window around the gene coding for the protein, extending 1Mb from the 5’ and 3’ untranslated regions (UTRs), based on the most extreme 5’ and 3’ ends of any transcript of the gene. We used 1Mbp as the approximate distance recently identified as the boundary between when a pQTL is more likely to be a *cis* rather than *trans* association^15^. Circulating protein level measurements were rank-inverse normalised at runtime, and age, age squared, sex, recruitment centre, 40 genetic principal components and Olink batch ID were included as covariates (**Methods**). In total, we tested 52,925,315 single (including single nucleotide variants and small insertions/deletions) and structural variant-protein associations. We identified independent variant associations by a combination of joint-modelling and forward-stepwise selection (**Methods**).

We annotated all genetic variants using Ensembl’s Variant Effect Predictor (VEP)^16^ (**Methods) and** used the output to categorise variants as gene-centric (e.g., coding, predicted intronic splicing, proximal-regulatory) and intergenic-regulatory (e.g., Ensembl regulatory regions, non-coding RNA) for aggregate-based association testing. Additionally, we performed aggregate testing on all non-coding (excluding proximal regions, to minimise overlap) variants in overlapping (1kbp overlap) 2kbp sliding windows. We additionally sub-categorised variants within a subset of aggregate units by measures of constraint (JARVIS^17^), conservation (GERP^18^) and/or predicted deleteriousness (CADD^19^). To identity independent rare non-coding genomic aggregate associations, we adjusted non-coding aggregate tests for common lead variants (MAF > 0.01), and all variants annotated as coding within the gene coding for the protein itself, which we henceforth refer to as the cognate gene, regardless of variant frequency. In total, we included 390,822,449 variants within our aggregate test analysis, with a mean of 179,319 per *cis* locus (approximately one variant per 10bp).

## Results

### We identified 1,425 rare variants associated with 1,450 circulating protein levels, with consistent effect estimates across multiple genetic ancestries

We identified 5,997 *cis* pQTL associations (MAC ≥ 5), 117 of which were structural variants (**Figs. 1&2; ST2)**. We identified at least one *cis* association for 1,126 proteins, with a median of four independent pQTLs per circulating protein. One hundred and seventy circulating proteins were associated with >10 independent pQTLs, including a maximum of 38 for *lair2.* The mean variance (within sample) explained jointly by all independent *cis* pQTLs for a given protein was 7.22%, with a median of 2.07%, similar to the estimates previously reported by Sun *et al*^14^. Of the 5,997 variant associations identified, 937 (15.9%) were in the rare frequency range (0.1% ≤ MAF < 1%), and 488 (8.3%) were very rare (MAF ≤ 0.1%). We refer to the 1,425 lead variants in both frequency bins as rare pQTL variants forthwith.

**Fig 1.**
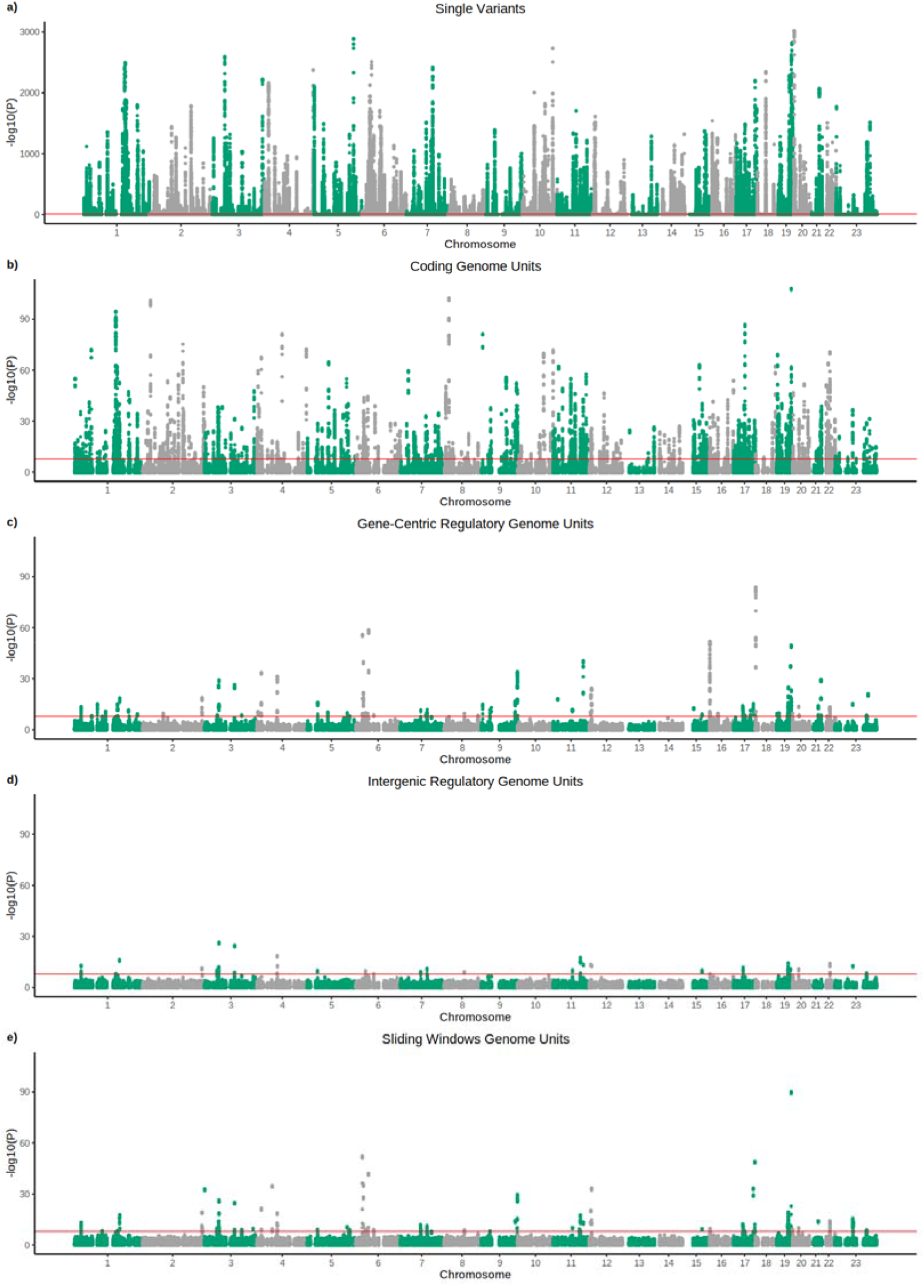
Manhattan plots showing associations between *cis* variants and regions with circulating protein levels, after adjusting for associated common variants and all coding variants of the cognate gene. The x-axis represents genomic position, and the y-axis shows – log10(p) for our *cis* results across all proteins, split into a) single variants, b) coding aggregates, c) gene-centric regulatory (proximal) aggregates, d) intergenic regulatory aggregates and e) sliding window aggregates. Red lines represent Bonferroni significance thresholds.

We additionally performed single variant testing for 430 and 451 individuals of genetically-inferred South Asian and African genetic ancestry respectively with both WGS and Olink proteomic data. Across the three genetically inferred genetic ancestries considered (European: EUR, South Asian: SAS and African: AFR), we observed a strong correlation of effect sizes for pQTLs in the EUR analysis between EUR and SAS individuals (r = 0.902), and weaker correlation between EUR and AFR individuals (0.646) and AFR and SAS individuals (0.645). Despite the much smaller sample sizes available for the SAS and AFR analyses each identified 100 independent pQTL variants (**ST3**), although power was limited to identify the full spectrum of pQTL variants across each *cis* locus.

We compared our single variant pQTL results with those of Eldjarn *et al* (2023)^12^, who analysed genomic data imputed from 150,119 UKB whole genome sequences in the 54,306 individuals with proteomic data. We found 2,586 of our 5,997 *cis-*pQTLs (42.8%) were in strong linkage-disequilibrium (r^2^ ≥ 0.8) with at least one of their signals for the same circulating protein. The overlap was larger (723 out of 1425; 50.7%) when considering only rare variants (MAF < 1%). There were 57 (3.93%) circulating proteins for which we identified more *cis*-pQTL variants than Eldjarn *et al*, and 1,057 (72.9%) proteins where they identified more variants. These differences may be partly driven by the sample size difference, and by differences in methods for conditional analysis: their analysis used forward stepwise conditional analysis to define conditionally independent pQTLs, while we performed both forward- and backward-conditional analysis steps implemented in GCTA CoJo^20^. It is likely that these differences in methodology have led to the differences in associated pQTLs between the two studies, highlighting the difficulties of interpreting multiple independent associated variants at the same locus.

### The majority of rare coding pQTLs are associated with reduced circulating protein levels

As an additional validation step, we tested whether coding variants within the cognate gene were associated with circulating levels of the protein. As expected, and consistent with the recent exome sequencing study^13^, 98.6% of pQTLs annotated as high-confidence loss-of-function as defined by LOFTEE^21^ were associated with reduced circulating protein levels (**ST2)**. The average effect of loss-of-function variants was -0.94 SD, equating to a reduction of raw circulating protein levels to approximately half (53.2%), with some notable exceptions (**Extended Table 1**). Consistent with variants in the last exon escaping nonsense mediated decay^22^, the estimated effects of loss-of-function variants were weaker towards the 3’ ends of the gene (**SF1**). Missense variants were associated with a much weaker effect, reducing circulating protein levels by 15.7% on average (**SF1)** and, as a negative control, the average effect of synonymous variants was close to zero.

**Table 1.** Genetic variants included in each grouping. UTR = Untranslated Region, 3’ = variants at the 3’ end of a transcript, 5’ = variants at the 5’ end of a transcript, GERP = Genomic Evolutionary Rate Profiling score (a measure of conservation), Start Gained/Lost = the inclusion or removal of a start codon, Downstream = downstream of a transcript, CADD = Combined Annotation Dependent Deletion score, AI = Splice AI (AI) score.

| CLASSIFICATION | MASK | CONSEQUENCES |
| --- | --- | --- |
| Proximal | 3' UTR | 3' UTR |
|  | 3' UTR (GERP>2) | 3' UTR (GERP>2) |
|  | 5' UTR | 5' Start Gained, 5' Start Lost, 5' Start Rest |
|  | 5' Start Gained | 5' Start Gained |
|  | 5' Start Lost | 5' Start Lost |
|  | Conserved and Intronic | Constrained |
|  | Downstream Any | Downstream |
|  | Downstream and conserved | Downstream with GERP>2 |
|  | Downstream and deleterious | Downstream with CADD>25 |
|  | Downstream and constrained | Downstream with JARVIS > 0.99 |
|  | Intron Splice Variant with AI>70 | Intron Splice Acceptor gain/loss with AI>70, Intron Splice Donor gain/loss with AI>70 |
|  | Splice Variant | Splice Region Variant |
|  | Upstream and conserved | Upstream Variant (GERP>2) |
|  | Upstream and deleterious | Upstream Variant (CADD > 25) |
|  | Upstream and constrained | Upstream Variant (JARVIS > 0.99) |
|  | Upstream Variant | Upstream Variant |
|  | RNA | Non-coding exon variant |
| Regulatory | Conserved, | Constrained and Conserved |
|  | Constrained and Intergenic |  |
|  | Conserved (GERP >2) Constrained and Intergenic | Constrained and conserved with GERP >2 |
|  | Regulatory Region Variant | Regulatory Region Variant |
|  | Conserved (phastCon 30) | Top 1% conserved variants in phastCon 30 window |
|  | Conserved (phastCon 100) | Top 1% conserved variants in phastCon 100 window |
|  | Phastcon100&30 and Conserved | Any phastcon variant (top 1%) for both phastcon 100 and 30 and conserved (GERP>2) |
|  | Phastcon100 and Conserved | Phastcon100 (top 1%) and conserved (GERP>2) |
|  | Phastcon30 and Conserved | Phastcon100 (top 1%) and conserved (GERP>2) |
|  | Phastcon100 and Conserved at any level | Any phastcon variant (top 1%) for phastcon 100 and conserved |
|  | Phastcon30 and Conserved at any level | Any phastcon variant (top 1%) for phastcon 100 and conserved |
|  | RNA | Non-coding exon variant |
| Coding | Synonymous | Synonymous |
|  | Missense | Missense |
|  | Missense with CADD>25 | Missense variant (CADD>25) |
|  | LoF | High Confidence Loss of Function |
|  | Splice Region | Splice Region Variant |
|  | Highly Damaging Splice Region | Splice region variant (spliceAI > 0.7) |

Relative to the background proportion, defined as the full set of coding variants tested, we observed an enrichment for rare pQTL variants annotated as loss-of-function (OR = 5.22 [4.24, 6.41], difference in proportion for rare pQTLs (ΔP) = 10.82%, Fisher’s exact *P*=1.16×10^-96^) and missense variants (OR = 2.19 [1.84, 2.60], ΔP = 17.6%, *P*=4.71×10^-77^). We additionally observed a depletion of splice-region (OR = 0.431 [0.294, 0.612], ΔP = - 7.60%, *P*=9.23×10^-36^) and synonymous (OR = 0.0958 [0.0616, 0.14], ΔP = -20.80%, *P*=8.39×10^-179^) variants.

We also performed aggregate-based association tests of variants within protein-coding genes. Aggregate based testing of rare coding variants identified 523 genes associated with circulating protein levels after adjusting for common (>0.1%) pQTLs from our single variant analysis (**ST4).** These genes were identified from 746 gene-protein associations, the majority based on including high-confidence loss-of-function variants alone (245; 32.8%), or all missense variants (291; 39%). Thirty-eight of these genes were not identified by the exome sequencing analyses performed by Dhindsa *et al* (2023) (**ST5**), potentially due to differences in sequencing coverage.

### The majority of rare non-coding pQTL variants are upstream of the cognate gene and are almost as likely to increase as decrease circulating protein levels

We identified 777 independent rare non-coding single variant-protein associations with one of 354 proteins (**Figs. 1&2; ST6)**. We stratified rare non-coding pQTLs into cognate and non-cognate groups based on annotation and most severe annotated consequence^23^, with priority given to annotations relative to the cognate gene. Of the 777, 551 (70.9%) were annotated as regulatory for either the cognate or another gene (**Figs. 2d and e**), and 226 (29.1%) were not assigned to a gene-centric or non-gene-centric annotation category (**Fig. 2e** “Unannotated”).

**Fig 2.**
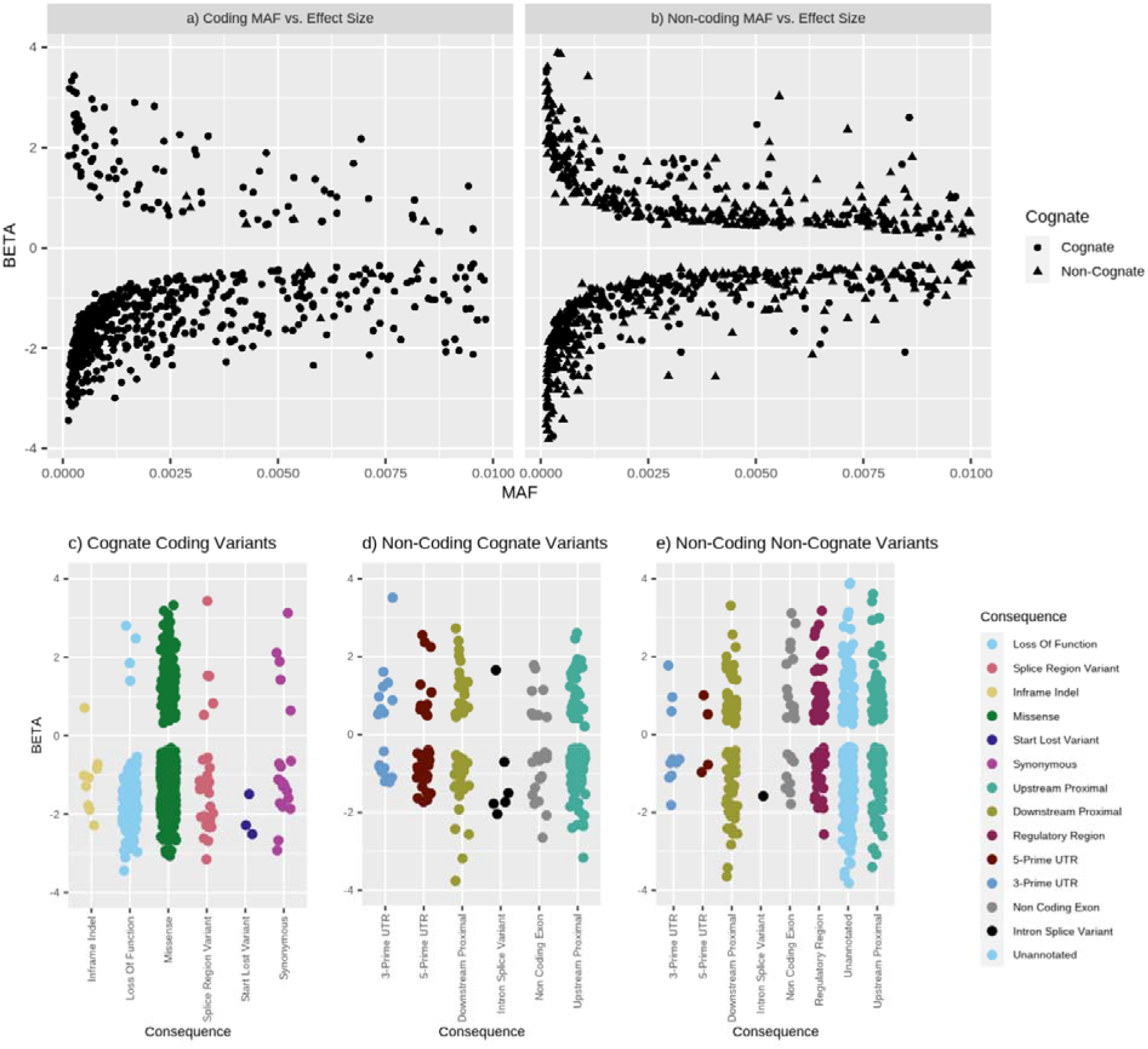
Effect size distributions of rare pQTL variants. Effect sizes for rare pQTL variants versus minor-allele-frequency for a) coding and b) non-coding pQTLs, and stratified by predicted consequence for c) coding variants in the cognate gene, d) cognate (variants annotated as regulatory for the protein-coding gene) non-cognate non-coding pQTLs.

These non-coding variants had an average absolute effect of 1.19SD (median 0.975SD), equating to 64.1% and 85.1% of the average absolute effect of rare loss of function and missense pQTLs on circulating protein levels respectively.

Rare non-coding pQTLs were distributed across the *cis* loci, with maximum distance from the cognate gene of 999 kb, and 247 of them (31.8%) were annotated as within a proximal or regulatory sequence of a neighbouring (non-cognate) gene but not the cognate gene itself (**SF2**). The single most strongly associated non-coding rare pQTL was located closest to, or in, the cognate gene 44.1% of the time.

Rare non-coding pQTLs were more evenly distributed between circulating protein increasing and decreasing effects (mean = -0.224, *P* sign = 8.08×10^-6^; **Fig 2d and e**), where ‘*P* sign’ is the p-value for a sign test, with effect sizes more balanced in the 551 annotated non-coding variants (mean = -0.162, *P* sign = 2.83×10^-3^; **Fig 2d and e**), compared to rare coding pQTLs (mean = -1.05SD, *P* sign = 8.77 ×10^-75^).

Non-coding rare pQTLs annotated as regulatory for the cognate gene (252; 32.4%) were more likely to occur in the upstream region of the gene than the downstream region. We observed that 120 (47.6% of 252) associations were annotated as upstream, 35 in the 5’UTR (13.9%), 6 (2.43%) predicted intronic splice acceptor/donor sites, 19 (7.69%) in the 3’ UTRs, 58 (23.0%) downstream and 14 (5.6%) in non-coding exons. The remaining 67.6% of signals were non-coding variants that were closer to, or resided within the intron of, a gene other than the cognate gene (**ST2**).

We then tested for enrichment withing different annotation categories for variants annotated as regulatory for the cognate gene (252/777). Based on the most severe predicted consequence for each individual variant tested in any *cis*-window, with consequences related to the cognate gene prioritised over consequences on other genes, we observed an enrichment for lead variants annotated in the 5’UTRs (difference in abundance between background and lead variants, ΔP = 11.9%, Fisher’s exact *P*=5.89×10^-19^), and intron splice sites (ΔP = 2.38%, *P*=1.53×10^-11^). We additionally observed a depletion of downstream proximal variants (ΔP = -15.3%, *P*=3.40×10^-7^) (**Fig. 3a**). We did not observe any evidence for enrichment for 3’UTR variants (*P* = 0.710), cumulatively suggesting that regulation of translation initiation is more important to protein levels than regulation of transcription termination or mRNA stability. We did not observe any significant enrichment when considering rare non-coding pQTLs that were annotated as regulatory for another gene (the non-cognate gene; 247/777) in the *cis* window, suggesting the majority of these pQTLs were unlikely to be operating through a separate gene, and are distal regulatory elements for the cognate gene **(Fig. 3b**).

**Fig 3.**
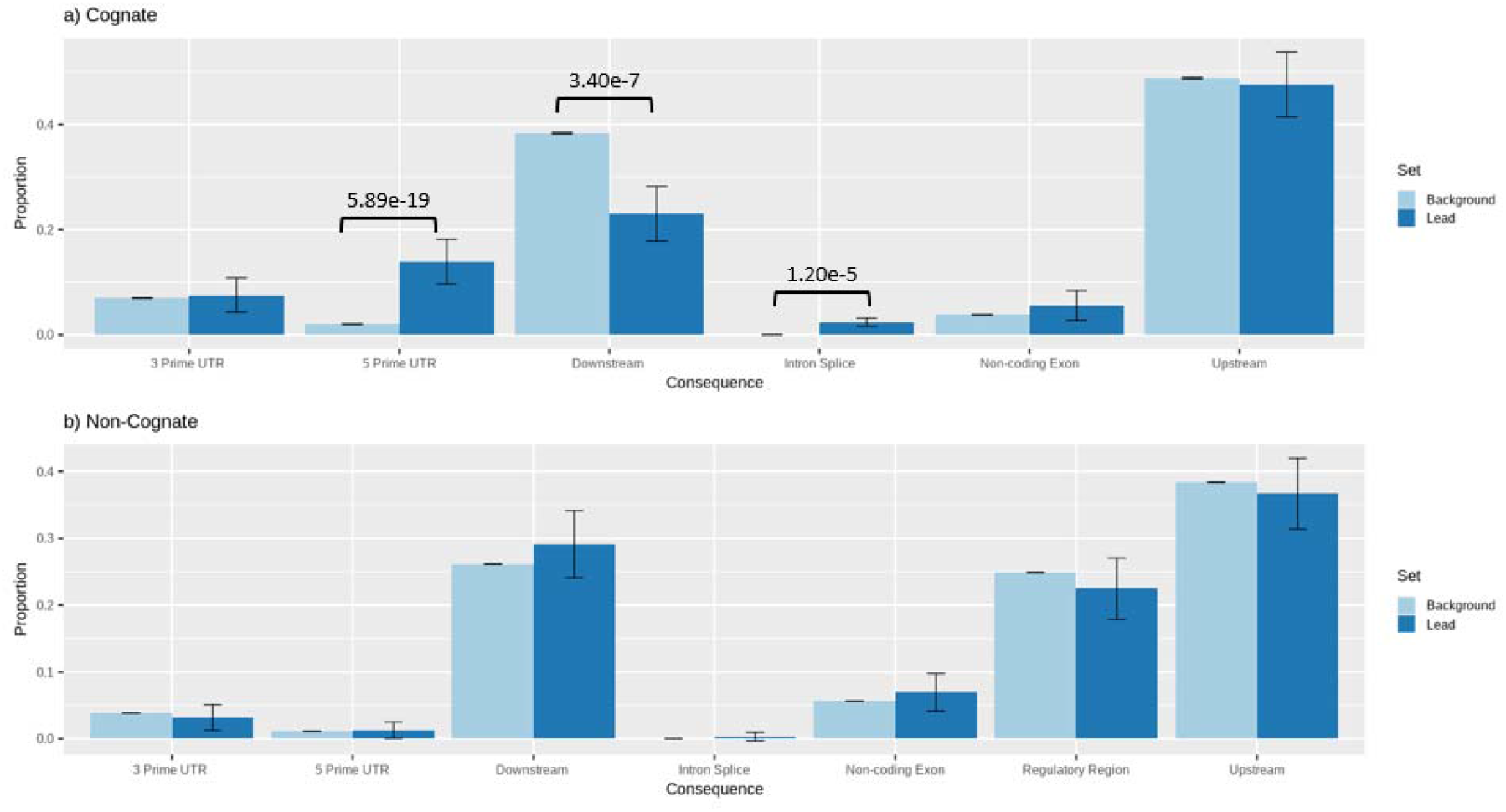
Distribution of annotations of lead pQTL non-coding variants compared to all variants tested. Proportion of variants in sets of lead variants (dark blue) compared to all variants tested (light blue), stratified by whether a variant was annotated to the cognate gene (a) or not (b). p-values are derived from Fischer’s exact test, only nominally significant (*P*<0.05) p-values are shown.

Given the challenges of grouping, analysing, and interpreting non-coding variants, we were interested in comparing the utility of several different computational metrics for variant prioritisation. Comparing the distribution of three widely-used measures of deleteriousness (CADD^19^), inter-species conservation (GERP^18^) and human variation constraint (JARVIS^17^) between non-coding pQTLs and all variants tested, we observed enrichment for deleteriousness (ΔCADD = 2.25%, *P =* 7.06×10^-10^), and constraint (ΔJARVIS = 8.89%, *P* = 2.96×10^-5^), but not conservation (**SF3)**. The observed enrichment was strengthened when considering only pQTLs annotated as regulatory for the cognate gene (ΔCADD = 4.97%, *P =* 2.60×10^-14^, ΔJARVIS = 17.4%, *P* = 3.92×10^-7^). Although we did not test the performance of other algorithms that aim to predict variant deleteriousness, most metrics - particularly of conservation - are primarily trained on the coding region, and better methods for variant prioritisation in the non-coding genome are urgently needed.

### A downstream variant produced the largest effect size observed for an annotated rare pQTL, and another single intronic ASGR1 variant was associated with 281 measured protein levels

Of the 777 rare non-coding variant-protein associations, 136 (17.5%) and 245 (31.5%) had absolute effect sizes larger than the average rare annotated pQTLs annotated as loss of function (mean absolute beta = 1.87SD) and missense (mean absolute beta = 1.41SD) respectively. The rare pQTL with the largest effect size was un-annotated and non-coding: 1:203482673:C:T, an intronic variant of *PRELP*, increased measured circulating levels of PRELP (beta = 3.89SD [4.47, 3.31SD], MAF = 2.85×10^-4^, *P* = 2.09×10^-46^). Further, the rare *annotated* pQTL with the largest effect size was also a non-coding variant: 19:51129025:IG:C:CTT, a downstream of *SIGLEC9*, which results in reduced measured levels of SIGLEC9 (beta = -3.75SD [-4.27, -3.23SD], MAF = 2.85×10^-4^, *P* = 2.09×10^-46^). In contrast, the largest observed coding effect was -3.44SD.

The 5’UTR variant with the largest change in protein levels was 18:597095:G:C, which increased circulating CLUL1 levels (beta = 2.55SD [2.22, 2.88SD], MAF=8.36×10^-4^, *P* = 5.29×10^-72^). This conserved position (GERP = 0.953 **- Methods)** impacts the 5’UTR of *CLUL1* (c.-48G>C, ENST00000420196.6 Clusterin-like protein 1), potentially creating a non-canonical (ACG) start site leading to a novel upstream open reading frame.

The 3’UTR variant with the largest effect size was 22:17110069:A:C, which increased circulating levels of IL7RA (beta = 3.51SD [2.60, 4.43SD], MAF = 1.30×10^-4^, *P* = 1.24 ×10^-14^) by impacting a conserved position (GERP = 5.06) in the 3’UTR of *IL7RA* (ENST00000319363.11:c.*249A>C).

Previous studies have shown that some *cis* pQTLs can have *trans* effects on multiple proteins. We therefore tested the association of each rare non-coding pQTL with each of the 1,450 measured protein levels. Based on the 1,424 identified rare single variants, we identified 677 additional *trans* variant-protein associations **(ST7)**. One rare small non-coding deletion, 17:7176936:CCCCCAGCCCCAG:C (MAF=0.8%), was associated in *cis* with circulating levels of two protein levels at the same locus (CLECL10A and TNFSR10B), and with 279 proteins in *trans*. This variant overlaps an Ensembl Candidate Cis-Regulatory Element (EH38E1844080) and lies within a third gene in the *cis* locus, in intron 4 of *ASGR1,* but showed no evidence of association with ASGR1 protein levels (*P* = 0.79). For all 281 (*cis* and *trans*) associations, the deletion was associated with *increased* circulating protein levels. Of the 281 proteins associated with this deletion, 275 (98%) were glycoproteins, representing a significant enrichment (binomial *P* < 2.2×10^-16^), in line with previous functional analysis^24^.

This *ASGR1* variant has also previously been associated with reduced risk of coronary artery disease, decreased LDL-C (LDL cholesterol), increased levels of alkaline phosphatase and vitamin B_12_^24^, and as a chronic inflammation marker^25^. In the UKB, we replicated the associations with LDL-C, HDL-C, alkaline phosphatase, and triglycerides (**ST8**). In an attempt to determine the causal pathway driving the association between the non-coding variant and LDL-C we examined the effect of an aggregate of loss-of-function variants (which would be predicted to *decrease* circulating protein levels) in each of the 275 associated glycoproteins on LDL-C. The strongest association of loss-of-function variants occurred with *GAS6* (Growth Arrest Specific 6, beta = 0.496 [0.305, 0.688], *P* = 3.67×10^-7^), suggesting that the variant may act partially through impacting *gas6* levels to reduce LDL levels.

### Aggregate testing identified 108 conditionally independent regulatory regions associated with bi-directional effects on circulating protein levels, after adjusting for common pQTLs and coding variation

Using aggregate variant tests, we identified 599 unique non-coding rare variant aggregates associated with one of 86 circulating proteins after adjusting for common single pQTLs and all protein-coding variants in the cognate gene **(ST9)**. After a further forward stepwise conditional analysis (**Methods**), 108 conditionally independent rare-variant aggregate non-coding regions remained (**ST10**). The conditionally independent non-coding aggregate-based tests used to identify these regions included 9 annotated to the 5’UTR (8.3%), 4 to the 3’UTR (3.7%), 31 annotated as gene-centric upstream (28.7%), 16 as gene-centric downstream (14.8%), 5 as predicted intronic splice acceptor/donor (4.6%), 6 as intergenic regulatory regions (5.6%), 8 as annotated to a non-coding RNA (7.41%) and 29 sliding windows agnostic to regional annotations (26.9%).

The majority, 55 (51%), of the 108 conditionally independent non-coding aggregate associations contained no individual rare pQTLs, suggesting those regions would not have been identified through single variant analysis. Of the remainder, 50 (46%) contained exactly one rare pQTL, and 3 (2.8%) contained more than one.

Five of the 108 (4.63%) conditionally independent non-coding aggregate associations were identified only when selecting highly conserved (GERP>2) variants, and 7 (6.5%) where identified when selecting highly constrained (JARVIS>0.99) variants. No aggregate associations were identified when selecting variants on predicted deleteriousness (CADD>25).

Different aggregate-based tests, before conditional analysis, often located the same noncoding region, with some identified by different sets of variants included in the test (e.g. conserved vs all variants), and some identified by overlapping 2kb sliding windows as well as annotated regions. For example, aggregate based testing of rare variants identified 12 non-contiguous regions across a 1.53Mb *cis* window associated with circulating levels of the interleukin receptor, IL17RB. Seven of these regions were each identified by pairs of 2kbp overlapping sliding windows. Two of these also contained regions annotated by Ensembl as regulatory which reached our association threshold (**ST9**). Conditional analysis collapsed these regions to two independent associations, one upstream of *CHDH* and RNA variants near *IL17RB*. In another example, aggregate based testing identified 20 non-contiguous regions across a 1.18 Mb *cis* region associated with circulating levels of the prostaglandin D synthase gene. These regions included 5 identified by sliding windows alone, and 15 as proximal to a gene. Our results thus highlight the need to perform conditional analysis for aggregate association testing, not only for single variants.

The vast majority of rare non-coding aggregate associations were identified by statistical tests that allow rare variants to be associated with both higher and lower trait values and that allow a large fraction of variants to be non-causal. Only three of the 108 (2.8%) rare non-coding conditionally independent aggregate regions were most strongly associated in a burden framework that assumes all rare variants result in effects in one direction. This is in strong contrast to coding-based aggregate tests, where 26.4% of unique aggregate tests were strongest in a burden framework. This difference suggests that rare variants in non-coding regions are likely to result in a mixture of trait increasing and trait decreasing effects, or that not all variants included in the aggregate test are causal, whereas a greater proportion of rare variants in coding regions are likely to be deleterious and causal.

For example, we found that an aggregate of rare non-coding variants in the 5’UTR of *CAPG* which each resulted in an additional start-site (5’ uATG gained) decreased circulating levels of CAPG (beta = -1.23SD [-1.61, -0.851SD], *P* = 2.06×10^-10^). As a second example, rare non-coding variants with a high (>0.7) SpliceAI score in the introns of *LRPAP1* decreased circulating LRPAP1 levels (beta = -1.39SD [-1.80, -0.983SD], *P* = 2.70 ×10^-11^).

When selecting only burden aggregates to identify aggregate associations (before performing conditional analysis within *cis* aggregates) **(ST11)**, we identified 37 significant unique aggregate associations with protein levels. Those aggregate tests altered measured protein levels by a mean absolute effect size of 1.35SD (signed mean -0.316). As with single variant rare pQTLs, the average effect of rare burden non-coding aggregates was considerably more balanced between circulating protein-increasing and decreasing effects, as compared to performing the same process for coding aggregates (signed mean = -1.33SD, *P* difference = 2.23 ×10^-4^; **ST12)**.

### Rare non-coding pQTL variants and aggregates were enriched in tissue-relevant panels and in proteins measurements with high cross-technology concordance

To determine the degree to which rare non-coding single variant and aggregate based pQTLs were present in tissue relevant non-coding regulatory regions, we performed two additional enrichment analyses. Firstly, we tested the hypothesis that rare non-coding pQTLs were more likely to be identified if the relevant protein was part of the cardiometabolic and inflammatory protein panels, rather than part of the neurology and oncology panels, on the basis that cardiometabolic and inflammatory processes are more relevant to circulating proteins. Secondly, we tested the hypothesis that rare non-coding pQTLs would be enriched in regions annotated as Ensembl regulatory elements in blood and liver cells ahead of 20 other tissue types (**Methods**) on the basis that blood and liver would be the tissue types most relevant to circulating proteins^26^. We further stratified these analyses by type of regulatory region – testing all regulatory regions, promoters, enhancers, transcription factor binding sites and CTCF binding sites.

Results of these tissue specific enrichment tests, separated by 22 tissue types and 4 panels, are presented in **Fig 4** (**ST13-18)**, and show that rare single variant and aggregate based non-coding pQTLs are enriched for proteins classified into tissue/process relevant panels. Furthermore, the results indicate that aggregate based tests, including those in region-agnostic sliding windows, identified more relevant regions compared to single variants, and pQTLs in regions outside of promoters were more panel specific than those in promotors. Whilst single variant pQTLs were enriched for proteins in all four panels, including the neurology and oncology panels (**Fig 4a**), aggregate pQTLs in predefined regulatory regions (**Fig 4b**) and sliding windows (**Fig 4c**), were enriched in inflammatory panels more than neurology and oncology panels. When limiting the analysis to promoters or enhancers we saw evidence for greater tissue and process specificity for enhancers. For example, single variant associations in promoters were enriched in all four panels (Fig 4d) but those in enhancers were only enriched in the cardio-metabolic panel (Fig 4g). Across the tissue types tested, we did not observe consistent enrichment for any specific tissue across the different analyses except in enhancers in the sliding-window analysis **(Fig 4i**) where the largest enrichment was seen for pancreas, vessel, liver, and heart tissues.

**Fig 4.**
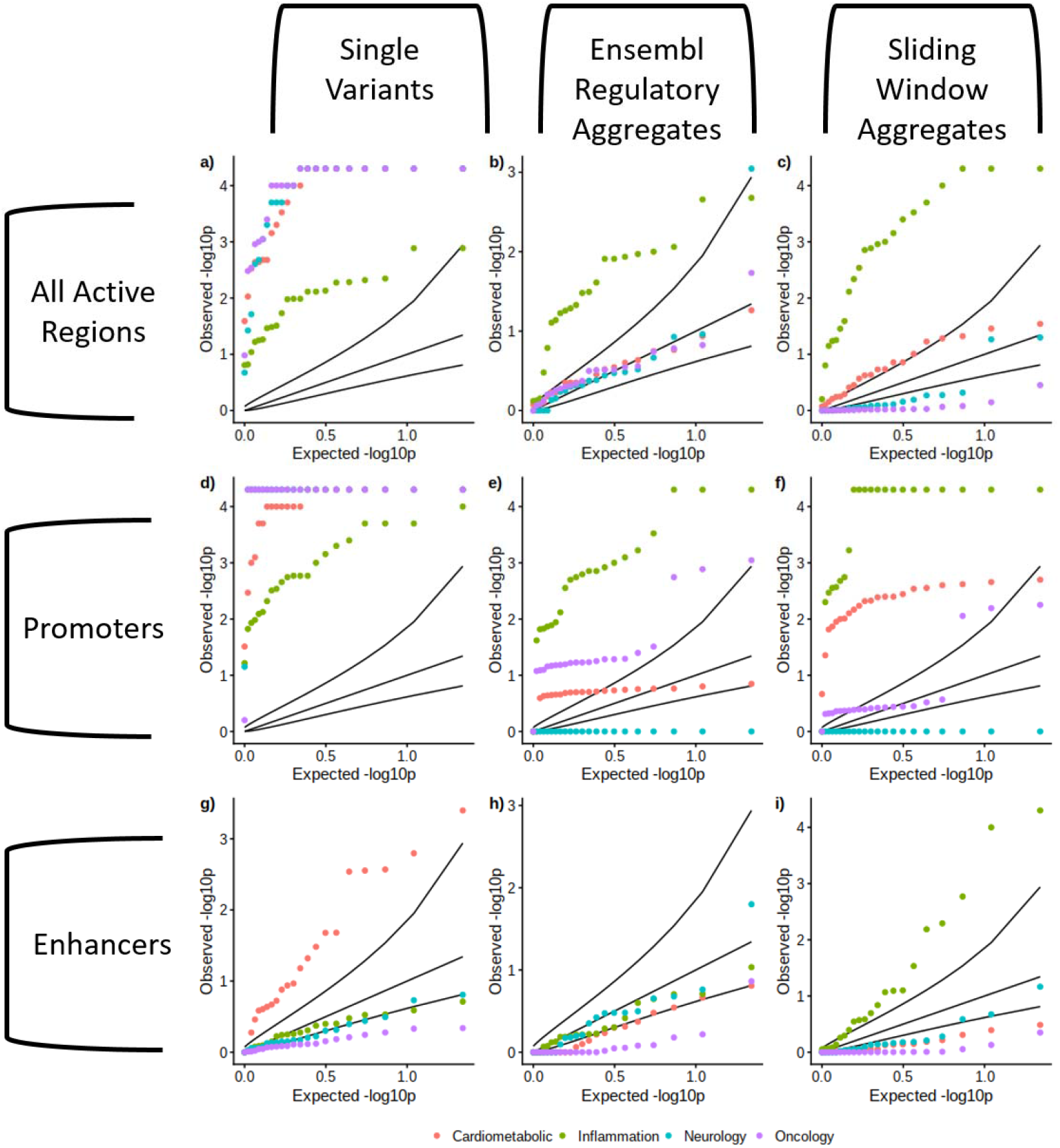
QQ plot for enrichment of loci within Ensembl predicted active regions within tissue groups. Empirical one-sided P-values for enrichment of signals within Ensembl predicted active regions within 22 tissue groups and four OLINK protein panels. Sub-figures a), d), and g) show enrichment for single variants, panels b), e), and h) show enrichment for Ensembl regulatory region based aggregate tests, and panels c), f), and i) show enrichment for sliding-window based aggregate tests. Panels a), to c) show enrichment within all predicted active regions, panels d) to f) for promoters, and g) to i) for enhancers.

To determine which circulating protein measures were consistent across different platforms, Eldjarn *et al* (2023)^12^ additionally compared the output of Olink (Explore 3072) technology with SomaScan v4 in 1,514 individuals of inferred Icelandic ancestry with whole-genome sequencing data. Based on the correlation between circulating protein measures across the platforms, and the similarity of lead *cis* pQTLs associated with the two measures, they identified 551 (out of 2,931) proteins as highly concordant (confidence tier 1 in their ST29). Of the 1,450 proteins (26.9%) in our study, 390 were highly concordant between the Olink and SomaScan technologies. Our pQTL associations, including those involving non-coding aggregate-based tests, were enriched in these 390 proteins. These 390 proteins represented 27% of the proteins we tested but included 37.8% (test of two proportions *P* = 1.45×10^-14^) of all our pQTLs and 42.2% of non-coding aggregate based pQTL associations (*P* = 1.31 ×10^-11^).

## Discussion

Using circulating protein levels as exemplar traits, we have shown that the analysis of WGS data enables the discovery of multiple rare non-coding variants and aggregates of rare variants associated with common phenotypes. Importantly, WGS data enabled us to consider more than six times the number of variants than would have been possible through single variant testing and identify associations between aggregates of rare variants and common phenotypes, using methods analogous to those used to aggregate coding variants in exome sequencing studies. The presence of multiple rare non-coding associations is consistent with the presence in the non-coding genome of most common variant associations identified by GWAS.

We have identified hundreds of novel non-coding rare aggregate and single variant associations with one of 1,450 measured protein levels in *cis*-windows 1Mbp either side of the cognate gene. We show that the effect sizes of non-coding associations can have similar absolute values compared to coding associations but are more balanced between circulating protein increasing and decreasing effects. We show that a single non-coding intronic deletion in *ASGR1* is associated with at least 281 distinct protein measurements, potentially by impacting glycoprotein turnover. Eldjarn *et al.* (2023)^12^ also identified 212 circulating proteins associated with this variant, and Dhindsa *et al.* (2023)^13^ identified 186 circulating proteins associated with coding variants in *ASGR1,* highlighting *ASGR1* as a potentially key regulator of glycoproteins.

We demonstrate that the 5’UTR and predicted intronic splice acceptor/donor sites were enriched for rare non-coding pQTL variants, whereas we observed a depletion of downstream genetic variants associated with pQTLs. Our results suggest that, where statistical power is limited, variant-association discovery could be prioritised in those regions where we observed an enrichment of association signals.

We additionally demonstrated the power of aggregate testing for non-coding regions, akin to work already done to aggregate functionally similar variants in coding exons. By testing rare genomic aggregates of non-coding elements, grouped by (for example) conservation, constraint, or predicted regulatory activity, or using sliding windows, we identified a further 55 conditionally independent regions of interest not identified by single variant testing alone.

Compared to aggregate based coding associations, non-coding genomic aggregate associations were enriched for regions with bi-directional effect and association tests that allow a large fraction of variants to be non-causal. This observation is consistent with the fact that prediction of variant effects and functional regions is less precise in the non-coding genome compared to the coding genome. However, the fact that we have identified non-coding associations with current annotations and data indicate that more discoveries in common phenotypes are likely as functional annotations improve and population genetic data accumulates.

We have also made some important advances for conditional analyses when considering non-coding aggregates. Due to the complex nature of linkage-disequilibrium (LD), it is extremely difficult to determine (without additional data or functional work) whether a coding signal is driving a non-coding signal, or vice-versa. To mitigate against this effect, we took a conservative approach and conditioned on all coding variants for the cognate gene. Where the cognate gene is not known, it may be necessary to condition on all coding variants within a pre-specified window determined by LD patterns.

There were a number of limitations to our study. First, we were not able to replicate our results in a separate study as we did not have access to similar data from other studies. However, a large proportion of our associations reached levels of statistical confidence far below our threshold. Furthermore, we observed an enrichment of associations for the 390 circulating proteins which showed high concordance between technologies, and effect sizes were consistent in the individuals of African and South Asian ancestry. Second, we cannot be certain that we have accounted for all possible sources of residual confounding by LD with coding or common variants. However, it is unlikely our associations are substantially affected by residual confounding from coding variants because they have very different features including the much more equal distribution between trait increasing and decreasing effects compared to coding associations. Third, the majority of our discovery analyses were limited to individuals of European ancestry because we only had access to both WGS and Olink data from 881 individuals of non-European ancestry. However, within the samples available, we did identify variants not detected in the individuals of European ancestry and observed strong correlation between effect sizes across the three main ancestries. Fourth, all circulating proteins were measured in blood. Although a large portion of tissue-specific proteins are only expressed in those specific tissues, we were limited to considering circulating protein levels. Finally, we limited our analysis to *cis* windows to limit the number of tests performed. There are likely to be many *trans* effects driven by rare non-coding variants that have not yet been detected.

In conclusion, using an exemplar, biologically proximal trait of circulating protein measurements, we have shown that there are likely to be a large number of rare non-coding variants with large effects on complex phenotypes waiting to be discovered.

## Methods

### UK Biobank and Whole Genome Sequencing

The whole genome sequencing performed for UKB had an average coverage of 32.5X, with a minimum of 23.5X, using Illumina NovaSeq sequencing machines provided by deCODE^27^.

The genome build used for sequencing was GRCh38: single variant nucleotide polymorphisms and short ‘indels’ were jointly called using GraphTyper^28^. deCODE found that the number of variants identified per individual was 40 times larger than that found using WES in the initial 150,000 release of whole genome sequences.

Of the 200,000 individuals whose genomes were sequenced, we found, using genetic principal components as previously described^29^, there were 183,803 individuals of European ancestry in this subset of the UK Biobank.

### Human Protein Expression Levels

Protein levels for 1,463 proteins for 54,304 UKB participants were profiled using Olink technology, as described in Sun *et al* 2023^14^, by the UK Biobank Pharma Proteomics Project. Quality control procedures were applied to the data before being made available for researcher use, including outlier removal etc. Protein levels were additionally log-2 transformed before release. After quality control filtering, 54,189 individuals with protein expression data were approved for analysis. Sun *et al* found no evidence of batch or plate confounding effects. Of the 10,248 genetic variants reported by Sun *et al*, we successfully lifted 10,243 to human genome build 38 using UCSC liftover^30^. In total, 10,193 (99.5%) of those genetic variants were also present in the UKB whole-genome sequencing data. Of that subset, 1,145 genetic variants previously identified lay within the *cis*-window considered here.

### Genetic Data Format

We performed a multi-allele splitting procedure on each of the 60,648 pVCF whole genome sequencing files provided by the UK Biobank using bcftools^31^ and then converted those pVCFs to plink^32^ (v2.0) pgen/var/fam format. All plink files which contributed to a chromosome were then merged to generate a single per-chromosome genotype file.

### Genetic Variant Exclusion

We excluded all variants from our association analyses if *GraphTyper*, the software used to by UK Biobank to perform genotype calling, assigned an *AAScore* which was less than 0.5^27^, denoting variant quality.

### Association Analyses

We performed both single variant and aggregate tests within *cis* loci for each of the 1,463 proteins measured in UKB. To define the *cis*-window, we first mapped each protein to a coding gene (see **ST1** for a small number of exclusions), and for each gene determined the longest transcript recorded by Ensembl. Based on the longest transcript, we then defined the *cis*-window as a 1Mb window either side of the 5’ and 3’ UTR of the transcript gene (limited by the beginning and ends of chromosomes), as well as the variants within the coding and intronic sequences. All association analyses were corrected for age, sex, age squared, UK Biobank recruitment centre (as a proxy for geography) the first forty genetic principal components, whole-genome sequencing batch and Olink plate.

#### Single Variant Association Testing

To identify *cis* single variants associated protein levels we first performed an association test for all genetic variants with a minor-allele-count of at least 5 using *regenie*^33^ (v3.14) in the *cis*-window. Lead variants were then selected in a conditional-joint analysis using *GCTA-CoJo*^20^ (diff-freq = 0.2, cojo-p = 1×10^-9^), with the UK Biobank whole-genome sequencing data, limited to individuals with proteomic data, as an LD reference panel.

If any lead conditionally-independent variant derived by *GCTA-CoJo* had an absolute joint-beta > 4 (determined by the limits of a normal distribution with 20,000 samples), we instead performed forward-stepwise conditioning. Forwards stepwise selection was performed by repeatedly performing association analyses at the single variant level: if any variant in each run was study-wide significant, we selected the variant with the highest p-value and reperformed association analyses conditional on that variant (and all previously selected variants).

#### Rare Variant Genomic Aggregate Testing

To identify non-coding, potentially regulatory regions of the genome which were insufficiently powered for single variant analysis, we subsequently performed non-coding rare-variant (minor allele frequency <0.1%) genomic aggregate association grouping variants according to proximal 5’, proximal 3’ or intronic. To test whether non-coding rare variant aggregate signals were caused by / confounded by residual LD and haplotype structure with common variants and or single variant signals we performed the following steps for each rare variant aggregate test result reaching Bonferroni p <0.05:

1. **To generate our primary non-coding discovery results** we adjusted for the common lead variants identified as independent signals in the joint (COJO) analysis (at MAF >0.1% ∼ MAC 40) AND adjusted for all genetic variants (regardless of p value) which we had annotated as coding in the gene which mapped to the protein of interest
2. **To identify independent non-coding aggregate associations,** we performed a forward stepwise regression. Starting from the most-strongly associated (genome-wide) non-coding aggregate (by p-value), per-protein, we perform an additional non-coding aggregate-testing run for any genome-wide significant aggregate adjusted for all variants in the top signal. This process is repeated, with more variants added, until no aggregate is genome-wide significant.
3. **To establish the extent to which our primary aggregate discovery results** could be due to a single low-frequency lead variant, we identified aggregate associations containing exactly one lead genetic variant.
4. **As a sensitivity step**, to establish the extent to which these results could be due to confounding linkage disequilibrium, we performed a further step where we adjusted for all pQTL single variants identified

Genome unit testing was performed for variants with a maximum allele frequency threshold of 0.1%, using *regenie,* based on the genetic units specified in Table 1. *regenie* performs four types of genome unit tests:

1. Standard BURDEN tests, under the assumption that each variant in a given gene unit mask has approximately the same effect size and sign on the phenotype
2. SKAT tests, where the sign of association of each variant in the unit is allowed to vary
3. ACAT tests, where the sign of association of each variant in the unit can differ, and only a small number of variants in the mask need be associated
4. ACAT-O, which is an omnibus test of BURDEN, SKAT and ACAT that aims to maximise the statistical power across the three tests

We performed each of the four statistical tests above for each mask for which a genome unit has at least one variant. Additionally, a singleton association test was performed for all variants with MAC=1 in each unit. *regenie* also estimated an ‘all-mask’ association strength for each genome unit, which is an aggregation of the test statistics of the individual masks. To ensure that this did not result in a mixing of non-coding and coding association statistics, we split each gene transcript into a coding transcript, which we tested for all coding masks, and a proximal transcript that we tested for all proximal masks. Regulatory genome units were either classified by their ENSR assignment, by the extent of a 1kb constrained window, or a phastCon conserved window. We named sliding windows masks by the region of the respective chromosome that they covered.

### Genetic Variant Annotation

We annotated all genetic variants using Variant Effect Predictor (VEP). Where possible, we assigned each variant to one of three *classifications*: coding, proximal-regulatory or intergenic-regulatory. A variant was classified as coding if it had an impact on an exon of **any** transcript; proximal-regulatory if the variant lay within a 5kbp window around a transcript or an intron, and was not already a coding variant in any transcript, and finally intergenic-regulatory if the variant fell within a conserved, constrained, non-coding exon region (details below), and was neither proximal or regulatory. We additionally tested variants in sliding windows of size 2000 base pairs, regardless of the number of variants in each window, with proximal and coding variants excluded to minimise hypothesis overlap.

We then assigned each variant to groupings, which we refer to as *masks*, according to their predicted consequence and location. We used five published variant scores to group variants by consequence:

1. **Genomic Evolutionary Rate Profiling (GERP)** The GERP score is a measure of conservation at the variant level^18^. We classified a variant as highly conserved if it had a GERP score >2.
2. **phastCon score** phastCon is a window-based measure of conservation across species^34^: either strictly mammalian (phastCon 30), or for all species (phast_100). We tested non-coding genome windows, i.e. excluding any window containing an exon, that had a phastCon score in the top percentile.
3. **Constrained Score** Constraint was calculated in windows of size 1kbp^6^ based on the local mutability and observed mutation rate of each window. We tested windows with a constraint z-score greater than or equal to four.
4. **SpliceAI score** The SpliceAI score^35^ is a measure of how well predicted each variant within a pre-mRNA region is of being a splice donor/acceptor, or neither. A variant was classified as a splice site with high confidence if it had an AI>70.
5. **Combined Annotation Dependent Deletion score (CADD)** The CADD score^19^ predicts how deleterious a variant is likely to be. We applied the CADD score only to coding variants and considered loss-of-function variants only if tagged as high confidence by VEP. Missense variants with CADD>25 were segregated for testing in a separate mask.
6. ***JARVIS* Score** The JARVIS score was derived to better prioritise non-coding genetic variation for association study, based on a machine learning model derived from measures of constraint^17^.

Each genome mask consisted of a number of variants with different *consequences*, based on their location, one of the above scores and/or predicted coding consequences. For example, for a variant to be classified as missense CADD>25, it must change a codon of an exon of a gene transcript and be predicted to be highly deleterious.

In Table 1 we present the full list of consequences assigned to each mask and classification.

We re-assigned variants that fulfilled two distinct criteria within a given genome unit to avoid duplication. In these cases, a variant was re-labelled as a combination of the two criteria and were assigned to any mask which selects variants from at least one of those criteria.

### Pseudo Genes

We assigned variants to pseudo gene transcripts if they contained pseudo-exons. However, pseudo exons **were not** excluded from proximal regions of non-pseudo gene associations, instead being tested as a regulatory genome unit. If a pseudo-exon overlapped with any significant genome unit signal, we performed a bespoke analysis.

### Heterogeneity Calculations

We used the R-package *metafor*^36^ to calculate all heterogeneity p-values between effect estimates, under the assumption of a fixed-effects model.

### ENSEMBL Regulatory Region Enrichment

We calculated the enrichment of overlap for both single variants and aggregate regions with ensemble regulatory regions, which are available for 118 tissues/cell-lines from ENSEMBL^23^. For each tissue, ENSEMBL additionally provide predictions on whether each region is active (or inactive, suppressed etc), and the type of regulatory activity (promoter, enhancer, CTCF binding site, TF binding site, open chromatin region). We subsequently exclusively considered regions that were predicted to be active, excluded cell-lines and cancer-derived tissues, and grouped the remaining tissue-types into 22 supergroups (see **ST9**). At the protein level, olink additionally grouped each protein into one of four panels (neurology, cardiometabolic, inflammation and oncology, as per the original publication^14^.

To determine the statistical enrichment, we performed bootstrapping over 10,000 simulations. For each simulation, we randomly selected a number of rare non-coding variants/aggregates determined by the set of our rare non-coding single-variants/aggregate tests from the *cis*-regions of the genome which we tested for association. We then determined the overlap of the randomly selected set of variants/aggregates with any of the regulatory regions (we re-performed this for each stratum of panel and tissue-type) and compared the distribution of the number of overlaps for any simulation with the number overlapping in our independent associations. We then assigned an empirical p-value to the observed overlap.

## Acknowledgements

This manuscript is part of the Stratification of Obesity Phenotypes to Optimize Future Obesity Therapy (SOPHIA) project. SOPHIA has received funding from the Innovative Medicines Initiative 2 Joint Undertaking under grant agreement No. 875534. This Joint Undertaking support from the European Union’s Horizon 2020 research and innovation program and EFPIA and T1D Exchange, JDRF, and Obesity Action Coalition www.imisophia.eu. GH has received funding from the Innovative Medicines Initiative 2 Joint Undertaking under grant agreement No 875534. ARW is supported by the Academy of Medical Sciences / the Wellcome Trust / the Government Department of Business, Energy and Industrial Strategy / the British Heart Foundation / Diabetes UK Springboard Award [SBF006\1134]. The research utilised data from the UK Biobank resource carried out under UK Biobank application number 9072. UK Biobank protocols were approved by the National Research Ethics Service Committee. TMF is supported by MRC awards MR/WO14548/1 and MR/T002239/1. The authors would like to acknowledge the use of the University of Exeter High-Performance Computing (HPC) facility in carrying out this work, funded by an MRC Clinical Research Infrastructure award (MRC Grant: MR/M008924/1). This study was supported by the National Institute for Health and Care Research Exeter Biomedical Research Centre. The views expressed are those of the authors and not necessarily those of the NIHR or the Department of Health and Social Care.

## Disclaimer

This communication reflects the author’s view: neither IMI nor the European Union, EFPIA, or any Associated Partners are responsible for any use that may be made of the information contained therein.

## Data Availability

Data cannot be shared publicly because of data availability and data return policies of the UK Biobank. Data are available from the UK Biobank for researchers who meet the criteria for access to datasets to UK Biobank (http://www.ukbiobank.ac.uk).

