## Supplementary Material: Extended Table 1 and Supplementary Figures 1-5 for "Whole genome sequencing analysis identifies rare, large-effect non-coding variants and regions associated with circulating protein levels"

Extended Table 1 & Supplementary Figures 1-5

*Rare Gain-of Function Loss-of-Function Variants*

We identified 4 rare variants which, despite being annotated as loss-of-function, were associated with increased circulating protein levels (**ET1**). The forty carriers of the splice region variant 8:23028330:C:T were unusually consistently affected, such that they generate a secondary, bimodal peak in raw *tnfrsf10b* measurement levels. On the raw scale, the splice variant was associated with more than 130x the mean protein levels (beta = 136.082 [122.428, 151.258], *P =* 3.39x10^-1804^), which was consistent across all carriers. The 1bp deletion was not associated with such dramatic changes on the raw scale (beta = 3.321 [2.80, 3.94], *P =* 1.023x10^-42^).

**Extended Table 1**: Four rare loss-of-function variants, as annotated by LoFTEE packaged in Variant Effect Predictor, which we estimated to have a gain-of-function effect on the protein produced by the cognate gene.

| **PROTEIN** | **VARIANT INFORMATION** | **EFFECT SIZE**  **[95% CI] (SD)** | ***P*** | **Biological Reasoning** |
| --- | --- | --- | --- | --- |
| *tnfrsf10b* | 8:23028330:C:T  ENST00000276431:c.748+1G>A  MAF = 9.69x10^-4^ | 2.80  [2.51, 3.10] | 1.04x10^-78^ | Both variants affect exon 5/9 of the transcript and would be expected to result in nonsense mediated decay; however, given the consistent and large effect size amongst all the carriers of the splice region variant, we can only hypothesise that they instead result in a protein product that is either more highly expressed in blood or more resistant to degradation. |
| *tnfrsf10b* | 8:23028480:GA:G  ENST00000276431:p.Ser200ProfsTer17  MAF = 3.77x10^-4^ | 2.48  [2.01, 2.95] | 4.75x10^-25^ |  |
| *nme3* | 16:1770682:C:T  ENST00000219302:p.Trp159Ter  MAF = 0.002883 | 1.85  [1.68, 2.03] | 8.67x10^-95^ | Likely that the transcript escapes NMD, resulting in a truncated protein that still includes the kinase domain but lacks the C-terminus, potentially reducing normal protein degradation mechanisms |
| *bst1* | 4:15737796:G:A  ENST00000514989:p.Trp95Ter  MAF = 0.005373 | 1.40  [1.32, 1.49] | 2.92x10^-233^ | This variant presumably causes the transcript to escape NMD, though we note that the variant also overlaps a putative enhancer (ENSR00001079930) downstream of the MANE Select transcript of BST1, which may also explain the association. |

**SF1.** Distribution of effect scores by relative gene positions for four primary coding consequence annotations. Stratified into high-confidence loss-of-function (HC LoF), low-confidence loss-of-function (LC LoF), missense and synonymous predicted consequences.

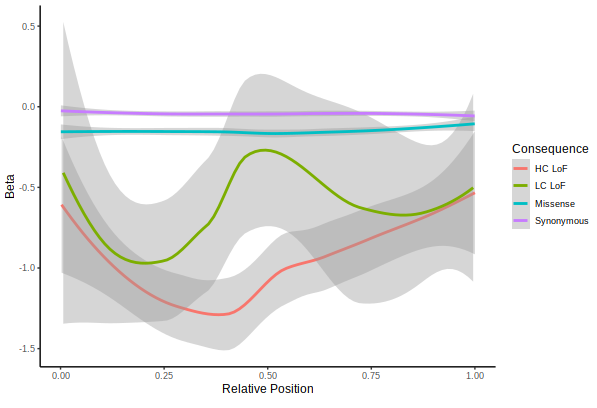

**SF2.** Associations between non-coding variants across all proteins; x-axis denotes relative distance from the gene, y-axis is the -log10p value. Variants between the 5’ and 3’ UTR were zeroed on the x-axis. Stratified by a) single variant associations and b) conditionally independent genomic aggregate regions, and c) shows the relative internal position of non-coding variants between the 5’ and 3’UTR of the longest transcript for each protein.

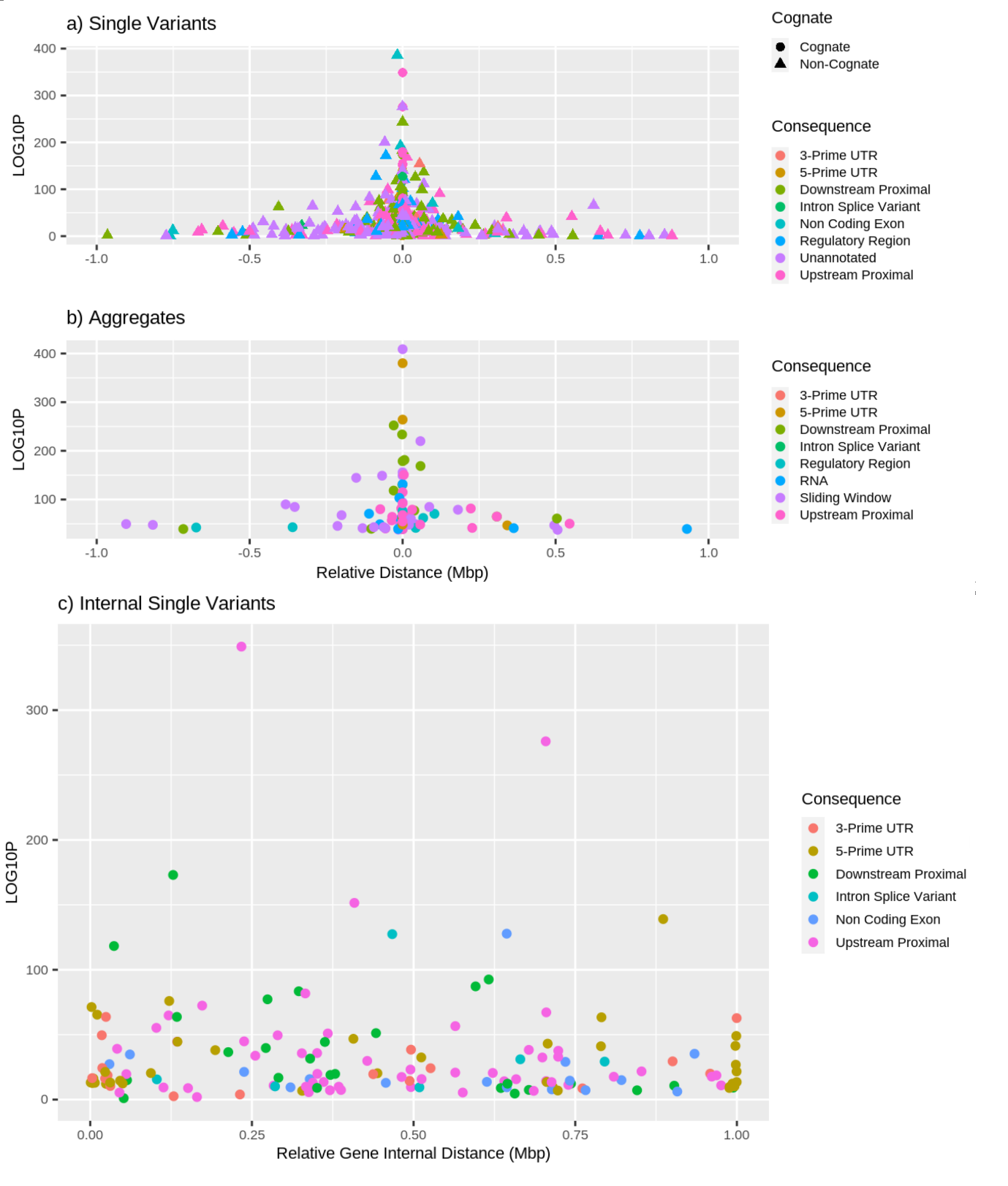

**SF3.** Variant scores CADD (deleteriousness), GERP (conservation) and JARVIS (constraint) for the lead pQTL variant at each *cis* locus.

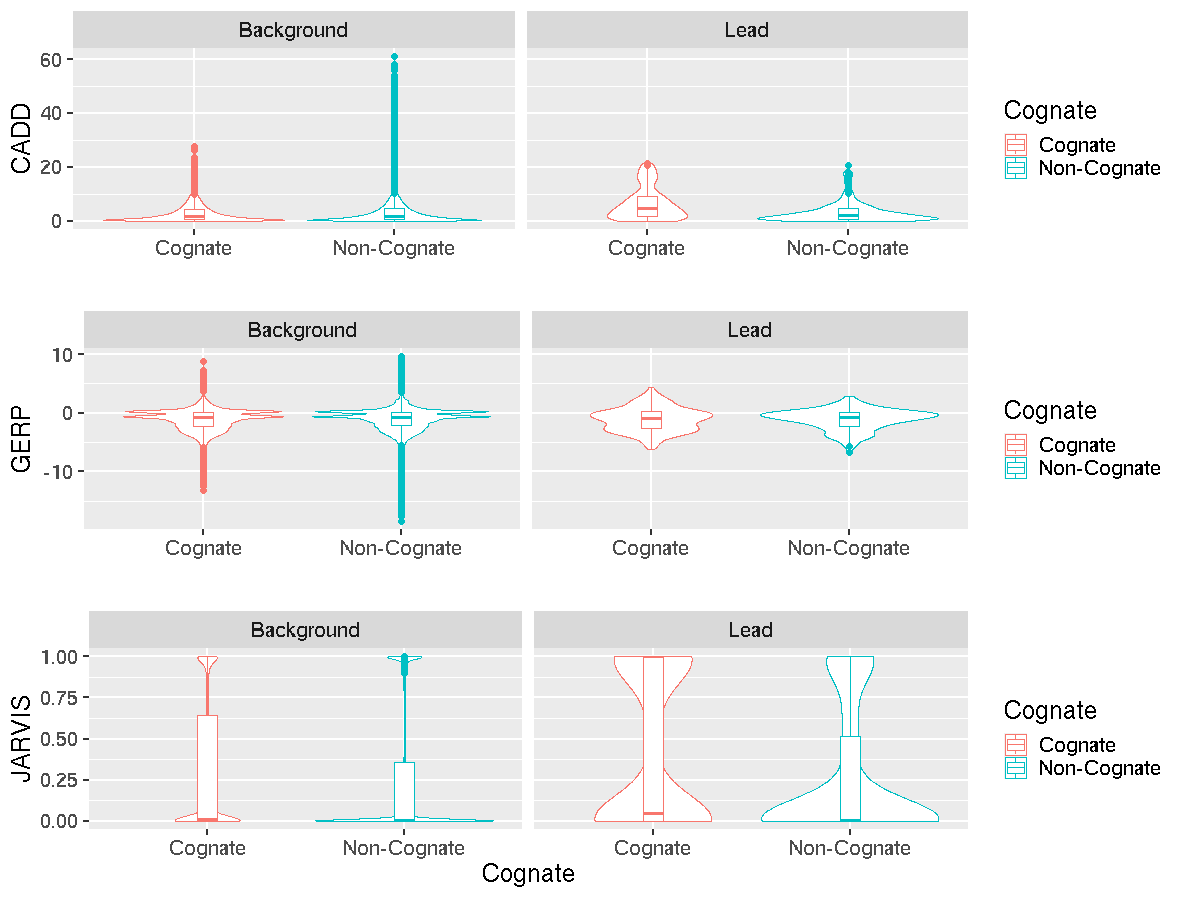

**SF4. QQ plot for enrichment of loci within Ensembl predicted active regions within tissue groups.** Empirical one-sided P-values for enrichment of signals within Ensembl predicted active regions within tissue groups. Panels a), e), i), m), q), and u) show enrichment for single variants, panels b), f), j), n), r), and v) show enrichment for aggregate tests, panels c), g), k), o), s), and w) show enrichment for sliding-window based aggregate tests, and panels d), h), l), p), t), and x) show enrichment for Ensembl regulatory region based aggregate tests. Panels a), to d) show enrichment within all predicted active regions, panels e) to h) show enrichment within CTCF binding sites, panels i) to l) for enhancers, m) to p) for open chromatin regions, q) to t) for promoters, and u) to x) for transcription factor binding sites.

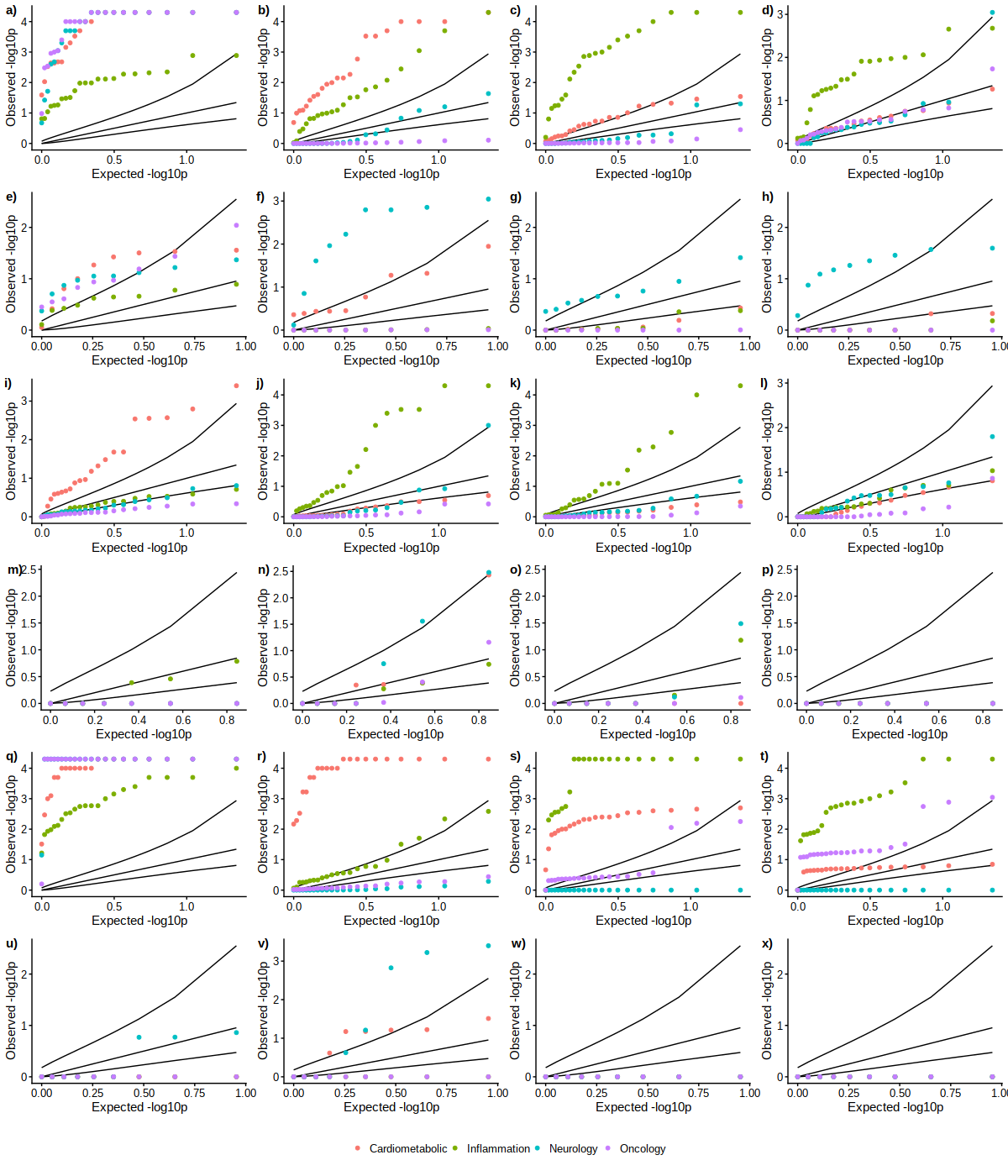

**SF5**. Proportion of Ensembl predicted active regions shared between blood and other tissues. Left panel: proportion of active regions overlapping between blood and other tissues as a proportion of the active regions for that tissue. Right panel: proportion of active regions overlapping between blood and other tissues as a proportion of the active regions in blood.

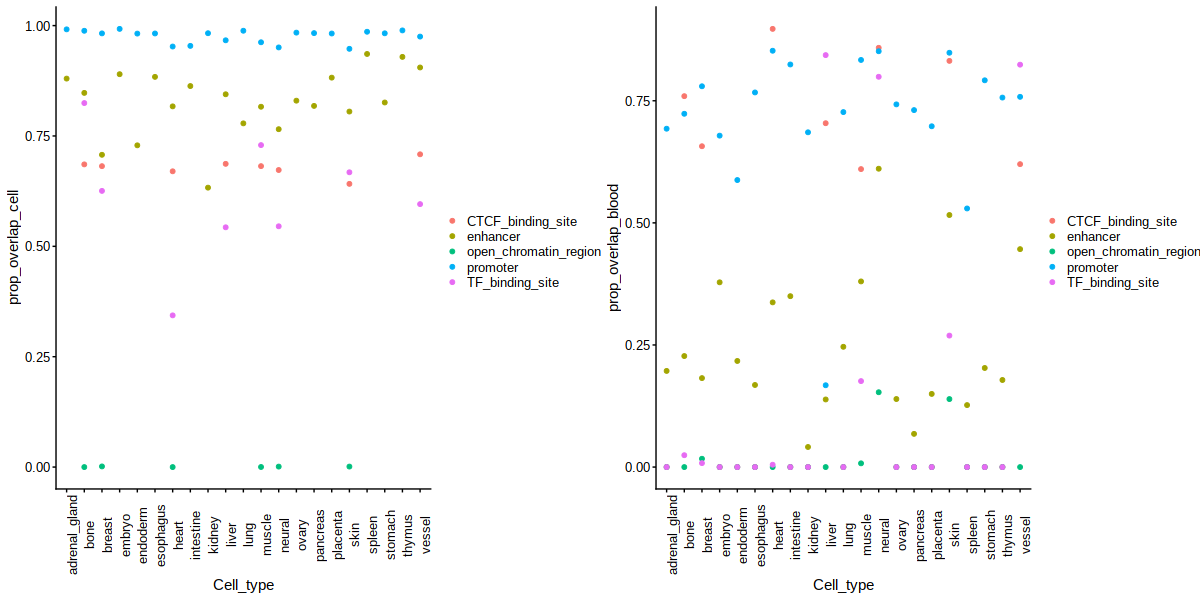
